# Modeling the circadian regulation of the immune system: sexually dimorphic effects of shift work

**DOI:** 10.1101/2020.11.30.403733

**Authors:** Stéphanie M.C. Abo, Anita T. Layton

## Abstract

The circadian clock exerts significance influence on the immune system and disruption of circadian rhythms has been linked to inflammatory pathologies. shift workers often experience circadian misalignment as their irregular work schedules disrupt the natural sleep-wake cycle, which in turn can cause serious health problems associated with alterations in genetic expressions of clock genes. In particular, shift work is associated with impairment in immune function, and those alterations are sex-specific. The goal of this study is to better understand the mechanisms that explain the weakened immune system in shift workers. To achieve that goal, we have constructed a mathematical model of the mammalian pulmonary circadian clock coupled to an acute inflammation model. shift work was simulated by an 8h-phase advance of the circadian system with sex-specific modulation of clock genes. The model reproduces the clock gene expression in the lung and the immune response to various doses of lipopolysaccharide (LPS). Under normal conditions, our model predicts that a host is more sensitive to LPS at CT12 versus CT0 due to a change in the dynamics of IL-10. We identify REV-ERB as a key modulator of IL-10 activity throughout the circadian day. The model also predicts a reversal of the times of lowest and highest sensitivity to LPS, with males and females exhibiting an exaggerated response to LPS at circadian time (CT) 0, which is countered by a blunted immune response at CT12. Overall, females produce fewer pro-inflammatory cytokines than males, but the extent of sequelae experienced by males and females varies across the circadian day. This model can serve as an essential component in an integrative model that will yield mechanistic understanding of how shift work-mediated circadian disruptions affect the inflammatory and other physiological responses.

**Author summary:** Shift work has a negative impact on health and can lead to chronic diseases and illnesses. Under regular work schedules, rest is a night time activity and work a daytime activity. Shift work relies on irregular work schedules which disrupt the natural sleep-wake cycle. This can in turn disrupt our biological clock, called the circadian clock, a network of molecular interactions generating biochemical oscillations with a near 24-hour period. Clock genes regulate cytokines before and during infection and immune agents can also impact the clock function. We provide a mathematical model of the circadian clock in the lung coupled to an acute inflammation model to study how the disruptive effect of shift work manifests itself in males and females during inflammation. Our results show that the extent of sequelae experienced by male and female mice depends on the time of infection. The goal of this study is to provide a mechanistic insight of the dynamics involved in the interplay between these two systems.

## Introduction

Most organisms from bacteria to humans are equipped with an internal biological clock, known as a circadian clock — a network of molecular interactions generating biochemical oscillations with a near 24-hour period [1]. In mammals, the circadian timing system consists of almost as many clocks as there are cells, as most cells house self-sustained and autonomous circadian oscillators [1]. This coordination of rhythms with the diurnal cycle is under the control of a central synchronizer, the suprachiasmatic nucleus (SCN), located in the ventral hypothalamus [2]. The SCN receives direct photic input from the retina, produces rhythmic outputs and orchestrates local clocks in the brain and peripheral clocks throughout the body [3].

Peripheral clocks can be coordinated by systemic cues emanating from the SCN [1], and they can be synchronized also by external cues such as temperature, feeding schedules and light [3]. In particular, the circadian circuitry in the lungs is exquisitely sensitive to environmental factors and exposomes [4], including air pollutants [5], cigarette smoke [6, 7], shift work [8–10], jet lag [11, 12], pathogens [13, 14] and much more. Of particular interest is the impact of circadian disruption on immune cell function, host defense and inflammation. The emerging picture is that the strength of the immune response varies throughout the day and that dysregulation of clock genes can lead to inflammatory disease or immunodeficiency [15].

Over the past decades, our societies have experienced rapid growth in the need for work in recurring periods other than traditional daytime periods. Research shows that shift work disrupts the natural sleep-wake cycle and feeding patterns [16], which may in turn cause serious health problems [17]. Here, we use mathematical modeling to study the effects of shift work, also known as chronic jet lag (CJL), on the lung circadian clock and consequently the immune response to inflammation. We address important questions: How do interactions between clock genes affect the strength of the inflammatory response at CT0 compared to CT12? Does the disruptive effect of shift work manifest itself differently in males and females? If so, what are the clock genes responsible for the sex-specific responses? Existing mathematical models of the circadian clock that focus on immunity can be classified into two categories: 1) models of the interplay between circadian rhythms and the immune system via neuroendocrine players (e.g. melatonin, cortisol) [18–20]; 2) models for the NF-*κ*B network modulated by the circadian clock [21]. The former do not model the core clock machinery, but rather use rhythmic hormones such as cortisol to drive the circadian variations in the system. The latter include the core clock system but with unidirectional coupling from the clock to the immune system. It is now known that the immune system can affect the circadian clock in a reciprocal manner [15, 22]. Given this observation, we have developed a model of the core circadian clock genes and proteins and their reciprocal interactions with the immune system under acute inflammation. Our mathematical model was extended to include the effect of shift work, represented as an 8h-advance of the circadian phase with sex-specific alterations in the expression of clock genes and proteins (see Fig 1).

**Fig 1.**
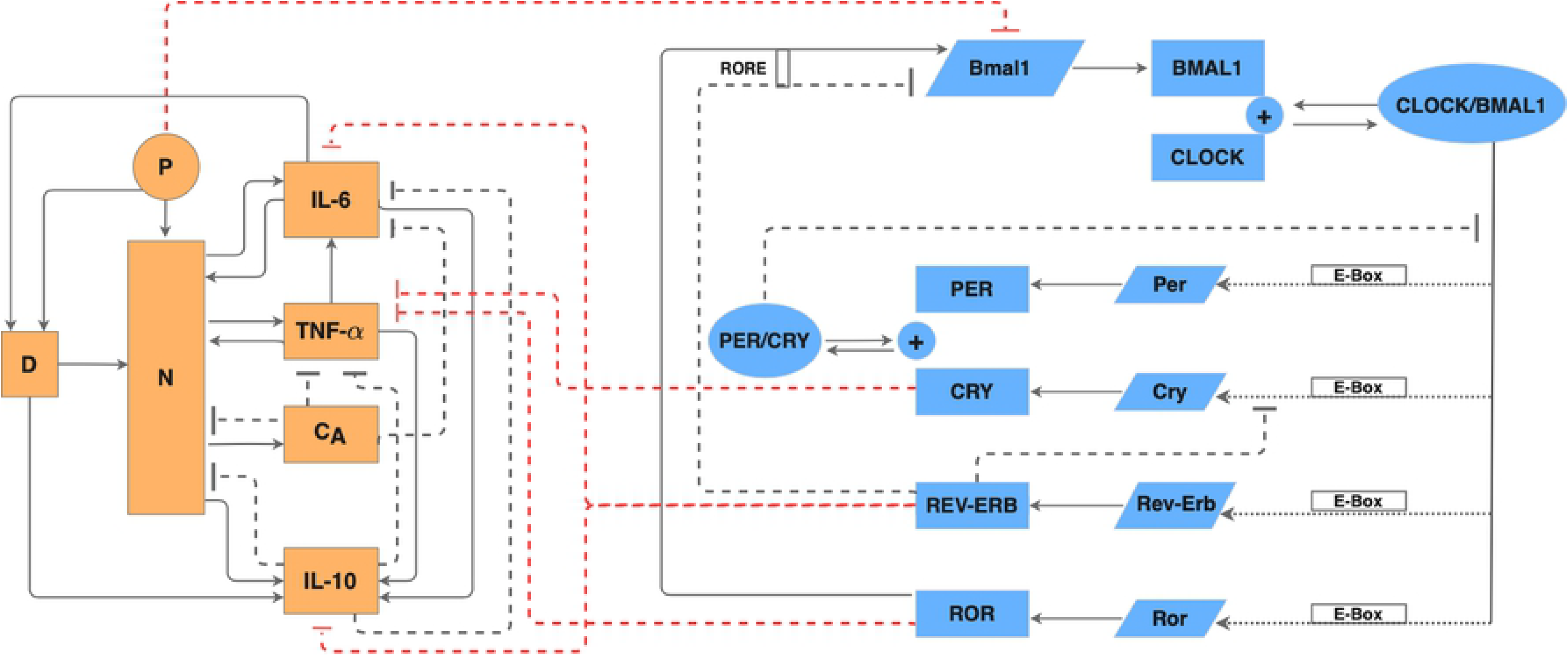
Regulatory network of the coupled immune system and circadian clock. Schematic diagram of the acute immune response model (orange shapes) and the circadian clock (blue shapes) in the lung of a rat. In the circadian clock model, slanted boxes denote mRNAs; blue rectangles denote proteins; ovals denote protein complexes. Dotted arrows represent transactivation; blunt dashed arrows represent inhibition. In the acute inflammation model, *P* denotes endotoxin; *D*, damage marker; *N*, activated phagocytic cells; *C*_*A*_, slow-acting anti-inflammatory cytokines.

The immune system is under control of the circadian clock. A primary means of circadian control over the immune system is through direct interactions of clock proteins with components of key inflammatory pathways such as members of the NF-*κ*B protein family [22]. This regulation is independent of transcription and allows the immune system to also reciprocally exert control over the function of the circadian clock [22]. Our model, which is composed of core clock genes (*Bmal1*, *Per*, *Cry*, *Rev-Erb* and *Ror*) and their related proteins as well as the regulatory mechanism of pro- and anti-inflammatory mediators (e.g. IL-6, TNF-*α* and IL-10), predicts temporal profiles of clock gene expression and cytokine expression during inflammation. This allows us to study how immune parameters respond to shift work-mediated circadian disruption. Moreover, we compare how the disruptive effect of shift work manifests itself differently in males and females.

## Materials and methods

We developed a mathematical model for simulating the circadian clock in the lung of a rat, the immune system under acute inflammation, and the interactions between the two systems. A schematic diagram that depicts the regulatory network is shown in Fig 1. Model equations and parameters can be found in Tables S1-S10 in supporting information.

### Circadian clock in the lung

The mammalian clock consists of interlocked transcriptional-translational feedback loops that drive the circadian oscillations of core clock components [23]. Both the master and peripheral clocks share essentially the same molecular architecture [24]. The activators CLOCK and BMAL1 dimerize to induce the transcription of target genes, including the *Period* genes (*Per1, 2, 3*), *Cryptochrome* genes (*Cry1, 2*), retinoic acid-related orphan receptor (*Rora, Rorb, Rorc*) and *Rev-Erb* nuclear orphan receptor (*Rev-Erbα, Rev-Erbβ*) to activate their transcription [1]. PERs and CRYs then heterodimerize and enter the nucleus to inhibit their own transcription by acting on the CLOCK-BMAL1 protein complex, and thus form the main feedback loop [1, 25, 26]. In the secondary loop, the nuclear receptors REV-ERB*α, β* and ROR*a, b, c* respectively repress and activate *Bmal1* transcription. The REV-ERBs, which also repress *Cry1* transcription [27], are essential for robust oscillations [28–30]. See the clock network in Fig 1, and refer to Refs. [3, 26, 28] for a detailed overview of the molecular clock architecture.

The present lung circadian clock model, inspired by Ref. [31], describes the time evolution of mRNA and corresponding protein concentrations of *Per*, *Cry*, *Rev-Erb*, *Ror*, and *Bmal1*. For simplicity, we grouped the three *Period* homologs (*Per1-3*) as a single *Per* gene and the two *cryptochromes* (*Cry1,2*) as a single *Cry* gene. Similarly, the two isoforms *Rev-Erbα* and *Rev-Erbβ* and three isoforms *Rora*, *Rorb* and *Rorc* are represented by single variables *Rev-Erb* and *Ror*, respectively. It was assumed that the CLOCK protein is constitutively expressed. We did not include post-translational protein modifications, considering that transport between the cytoplasm and the nucleus is rapid on a circadian timescale [32]. The time evolution of the core clock genes and proteins is described by Eqs. 1–12 in Table S2.

### Acute immune response

The acute immune response model, inspired by Refs. [33] and [34] with some modifications, consists of eight variables: endotoxin concentration (*P*); the total number of activated phagocytic cells (*N*, which includes activated immune response cells such as neutrophils and monocytes); a non-accessible tissue damage marker (*D*); concentrations of pro- and anti-inflammatory cytokines, namely IL-6, TNF-*α* and IL-10; a tissue driven non-accessible IL-10 promoter (*Y*_*IL*10_); and a state representing the level of slow acting anti-inflammatory mediators (*C*_*A*_), which comprises slow-acting anti-inflammatory agents such as cortisol and TGF-*β*1.

The introduction of bacterial insult in the system activates the phagocytic cells, *N*, and inflicts direct tissue damage, *D* [35]. This is different from the work of Roy et al. [33] in which endotoxin only activates phagocytic cells. The activated cells up-regulate the production of inflammatory agents (TNF-*α*, IL-6, IL-10, and C_*A*_) [36]. The pro-inflammatory cytokines TNF-*α* and IL-6 exert a positive feedback on the system by further activating *N*, as well as up-regulating other cytokines [36, 37]. The anti-inflammatory cytokines IL-10 and C_*A*_, on the other hand, have a negative feedback on the system. They inhibit the activation of *N* and other cytokines [38, 39]. In our model, *D* contributes to the up-regulation of IL-10 and is up-regulated by IL-6 because it has been shown that IL-6 is associated with the development of sepsis [40–42]. This differs from Ref. [33] in which damage is up-regulated by *N*. Note also that *D* should not be interpreted directly as a cell type in the model. The acute inflammatory response is described by Eqs. 13–20 in Table S2; see also Fig 1.

### Coupling between the circadian clock and the immune system

Most studies on circadian-immune interactions have focused on *Bmal1*, since inactivation of this gene is a convenient way to abrogate clock function [13, 43, 44]. Thus, care should be taken in distinguishing *Bmal1*-specific effects from downstream effects because other clock components act as intermediaries. The inhibitory effects that the circadian clock and the inflammatory response have on each other are shown in Fig 1. Specifically,

- **CRY proteins.** *Cry1*^−/−^*Cry2*^−/−^ mice exhibit an elevated number of T cells in the spleen with increased TNF-*α* levels [22, 45]. Other studies showed that *Cry1* and *Cry2* double KO in fibroblasts and bone-marrow-derived macrophages (BMDMs) leads to increased *Il6* and *Tnf-α* mRNA and an hypersensitivity to lipopolysaccharide (LPS) infection [15, 46]. Furthermore, the NF-*κ*B signaling pathway was shown to be constitutively activated in *Cry1*^−/−^*Cry2*^−/−^ BMDMs [46]. Due to the ensuing higher constitutive inflammatory state, *Cry1*^−/−^*Cry2*^−/−^ mice exhibit increased infiltration of leukocytes in lungs and kidneys [47]. Therefore, CRYs play an important anti-inflammatory role by downregulating inflammatory cytokines. Because IL-6 is inducible with TNF-*α*, effects on IL-6 in CRY double KO experiments are primarily mediated by TNF-*α* [46]. In our model, CRY directly inhibits the production of TNF-*α*, hence indirectly inhibits the TNF-*α*-induced IL-6 production (Eq. 17 in Table S2).
- **ROR proteins.** Similarly to the CRY proteins, experiments have shown that Ror*a*^−/−^ mice, also known as the staggerer mutant, exhibit higher levels of IL-6 in bronchoalveolar lavage fluid, which renders them more susceptible to LPS lethality [48]. Interestingly, staggerer mutant mice have an increased production of IL-6 and TNF-*α* in mast cells and macrophages after LPS stimulation [49, 50]. Furthermore, overexpression of ROR*a* in human primary smooth muscle cells inhibits TNF-*α*-induced expression of IL-6 [51]. The present model assumes that ROR downregulates TNF-*α*, thus indirectly downregulates IL-6 (Eq. 17, Table S2). Recall that the model does not distinguish between the three isoforms *Rora*, *Rorb* and *Rorc*.
- **REV-ERB proteins.** There is compelling evidence for a role for REV-ERB*α* in the control of the immune system. REV-ERB*α* is encoded by *Nr1d1* and *in vivo* challenge of *Nr1d1*^−/−^ mice with LPS leads to IL-6 upregulation in serum in comparison to wildtype animals [52]. REV-ERB*α* represses *Il6* expression not only indirectly through an NF-*κ*B binding motif but also directly through a REV-ERB*α* binding motif in the murine *Il6* promoter region [53]. A more recent study showed that the dual mutation of REV-ERB*α* and its paralog REV-ERB*β* in bronchial epithelial cells further augmented inflammatory responses and chemokine activation [54]. REV-ERB*α* also negatively affects the expression of anti-inflammatory cytokine IL-10. *Rev-Erbα* mRNA binds to the IL-10 proximal promoter and represses expression in human macrophages [55]. Together, these studies reveal the role of REV-ERB*α* as an equilibrist. In our model, REV-ERB directly inhibits the production of IL-6 and IL-10 (Eqs. 16 and 18 in Table S2, respectively). We note that the two isoforms *Rev-Erbα* and *Rev-Erbβ* are represented by a single model variable *Rev-Erb*.
- **Inflammation.** In a reciprocal manner, inflammation induced by agents such as LPS, TNF-*α*, and IFN-*γ* [56–60] or acute bacterial infection [61] can affect the circadian clock. In particular, rodent studies indicate that LPS transiently suppresses clock gene expression and oscillations in the SCN and peripheral tissues [32, 60, 62, 63]; notably, a number studies show significant suppression of *Bmal1* [15, 62, 64]. The inhibition of the circadian mechanism during endotoxemia lasts for at least 24 h [60, 64]. To represent the sustained effect of a bacterial infection on the circadian clock, we introduced a filter function for LPS (Eq. 21 in Table S2), which acts on the clock through its inhibition of *Bmal1* (Eq. 5 in Table S2). The filter function decays linearly over 24h and causes circadian disruption for at least this amount of time. We assume that the effects of cytokines such as TNF-*α* are incorporated in the net effect of LPS on clock genes, and so we do not include direct links from cytokines to clock genes and proteins.

### Model parameters

Most of the model parameters are not well characterized, and were estimated by fitting model dynamics to experimental data. This is done in a two-step process: we first fit the circadian clock model in isolation, in an infection-free state. This is done using data on the expression of circadian genes in the mouse lung under a constant darkness regime (*CircaDB* : http://circadb.hogeneschlab.org). In a second step, we fit the acute inflammation model together with the clock-inflammation coupling, without changing the circadian clock parameters. This is done by simultaneously fitting experimental measurements of the cytokines IL-6, TNF-*α*, and IL-10 in rat following the administration of endotoxin at 3 mg/kg and also at 12 mg/kg [33, 34]. This fitting was conducted with the coupling with the clock model taken into account.

Model parameters are shown in Tables S3–S10. It is noteworthy that the present model is based on the rat, whereas the parameters for the circadian clock and for the clock-inflammation coupling were based on mouse data. However, while species differences exist, core clock gene expressions of the mouse and rat lungs exhibit substantial similarities [65].

### Sexual dimorphism in clock gene alterations under circadian disruption

In their 2012 study, Hadden et al. reported sexual dimorphism in clock gene expression in the lungs of mice exposed to chronic jet lag [12]. Male and female mice were assigned to either remain in a LD12:12 regimen or to undergo experimental chronic jet lag (CJL). Under the CJL regimen, mice were subjected to serial 8-h advances of the light/dark cycle every 2 days for 4 weeks. Then using quantitative Polymerase chain reaction (PCR) to measure the relative amount of clock gene mRNAs, Hadden et al. observed that *Rev-Erbα* gene expression is upregulated in CJL males and downregulated in CJL females by 98% and 70% on average, respectively. *Bmal1* is downregulated in CJL females only by 43% on average, while *Clock*, which forms a heterodimer with *Bmal1* (CLOCK:BMAL1) [66], is downregulated in males only by 26%. The repressors *Per2* and *Cry2* are both upregulated when compared with same-sex control animals, although *Cry2* upregulation was not significant for CJL males. In particular, *Per2* and *Cry2* increased by 497% and 69%, respectively in CJL female mice, while *Per2* increased by 230% in CJL male mice. The authors did not test the effects of chronic jet lag on *Ror* gene expression. This could be explained by the fact that *Ror* is not directly related to the shift work phenotype. Indeed, the association between *Ror* and shift work disorder has been shown to be weak at best [67].

We used this information to create separate mathematical models of the lung circadian clock for males and females undergoing CJL. The decrease in *Clock* mRNA for CJL males is not taken into account because we do not model this gene explicitly and only constitutively represent the associated CLOCK protein. In addition, NPAS2, a paralog of CLOCK, has been shown to compensate for the loss of CLOCK in peripheral circadian oscillators [68, 69]. We note also that the baseline immune system likely differs between the sexes, but due to insufficient quantitative data, we were only able to construct one baseline model. Figure 2 shows the relative abundance of clock genes during CJL, normalized by CJL male values.

**Fig 2.**
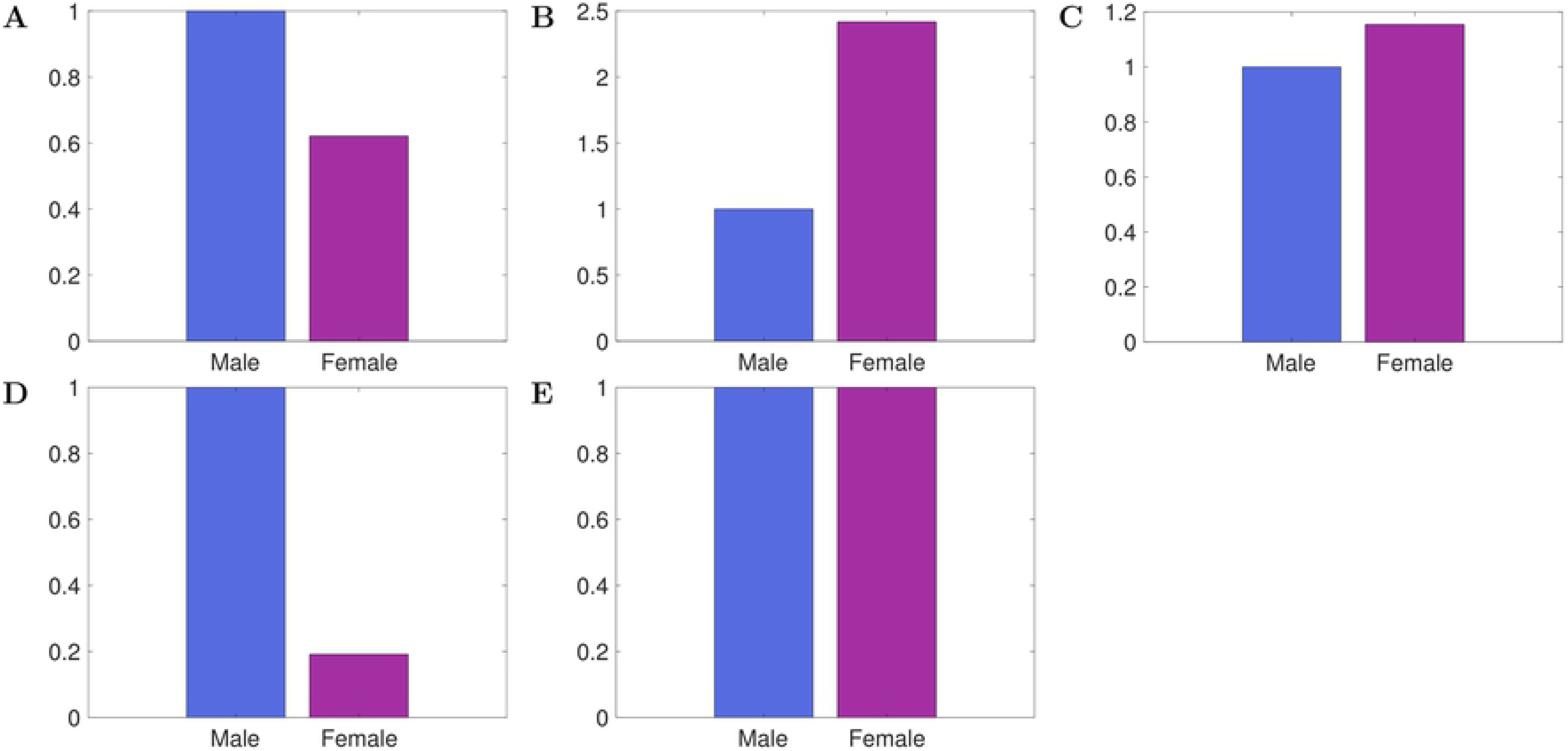
Relative abundance of mRNAs for CJL males and females, normalized by CJL male values. *Bmal1* (A), *Per* (B), *Cry* (C), *Rev-Erb* (D) and *Ror* (E).

## Results

### Expression of clock genes in lungs is accurately reproduced by the model

Using the baseline model parameters (Tables S3-S10), the circadian clock model predicts limit-cycle oscillations in the expression levels of all clock components with a system period of 24 h. Figure 3 compares the predicted time profiles of *Bmal1*, *Per*, *Cry*, *Rev-Erb* and *Ror* with experimental observations for *Bmal1*, *Per2*, *Cry1*, *Rev-Erbα* and *Rorc*. We refer to the onset of the rest phase of night-active organisms as circadian time (CT) 0 and to the onset of activity as CT12 [72]. These two times correspond to the onset of the light and dark phases, respectively. Good agreement can be observed between the predicted and experimental temporal profiles.

**Fig 3.**
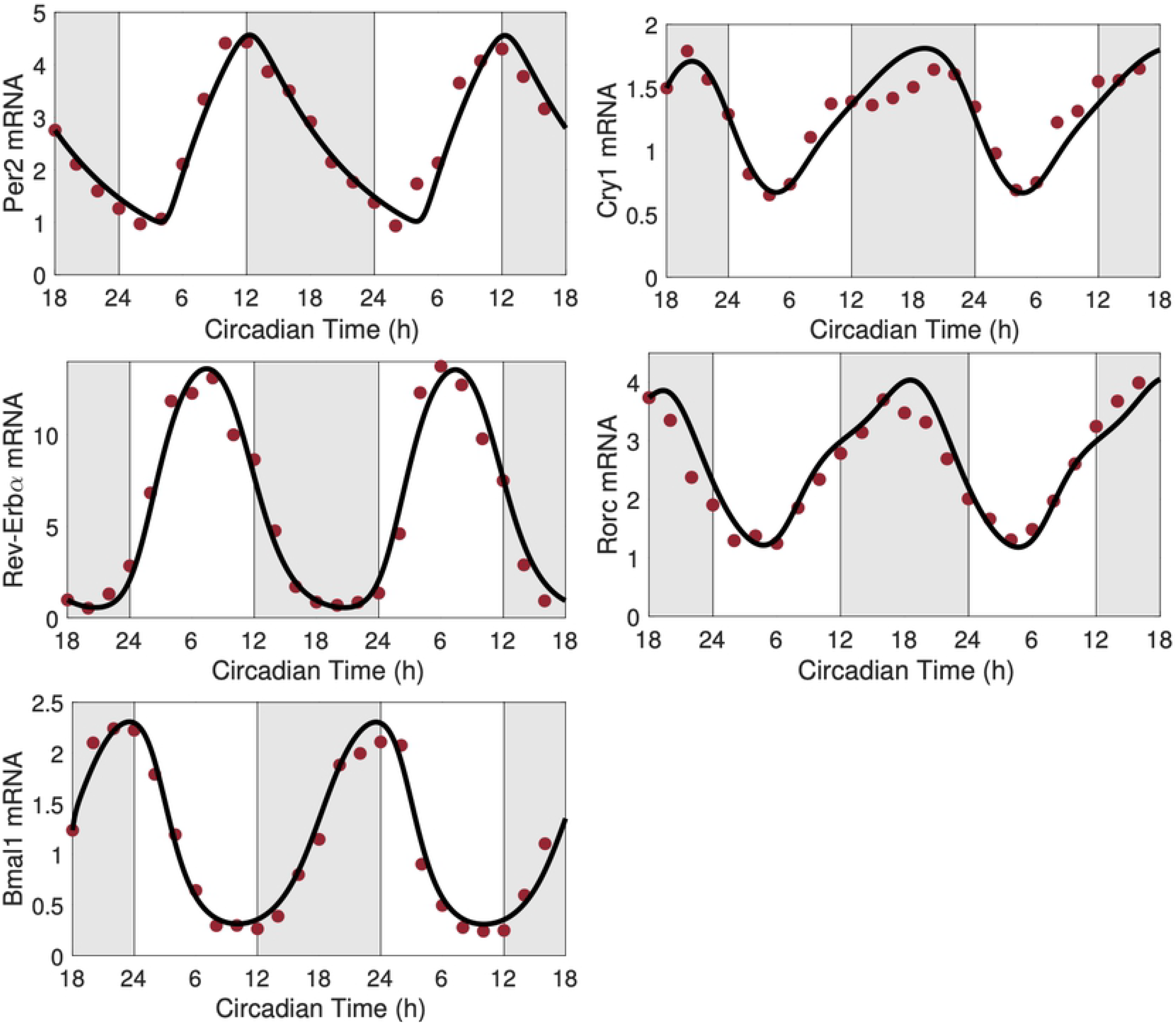
Predicted clock gene time profiles. Comparison of predicted time profiles (solid lines) for *Per*, *Cry*, *Rev-Erb*, *Ror* and *Bmal1*, with experimental data (circles) for *Per2*, *Cry1*, *Rev-Erbα*, *Rorc*, and *Bmal1* mRNA expression levels obtained in mouse lungs in constant darkness. Gray shading and white regions correspond to activity and restcycles, respectively.

### Cytokine dynamics during endotoxemia are accurately reproduced by the model

We estimated the parameter values for the acute inflammation model by simultaneously fitting experimental measurements of the cytokines IL-6, TNF-*α*, and IL-10 at endotoxin doses 3mg/kg and 12mg/kg. The predicted time profiles are shown in Fig 4, left and center columns. The model performs similarly well for all three cytokines, and is able to capture the second peak in IL-10 expression. We note that the predicted IL-10 concentrations at 1 h and after 10 h are slightly underestimated compared to the value recorded experimentally for the endotoxin dose 12mg/kg.

**Fig 4.**
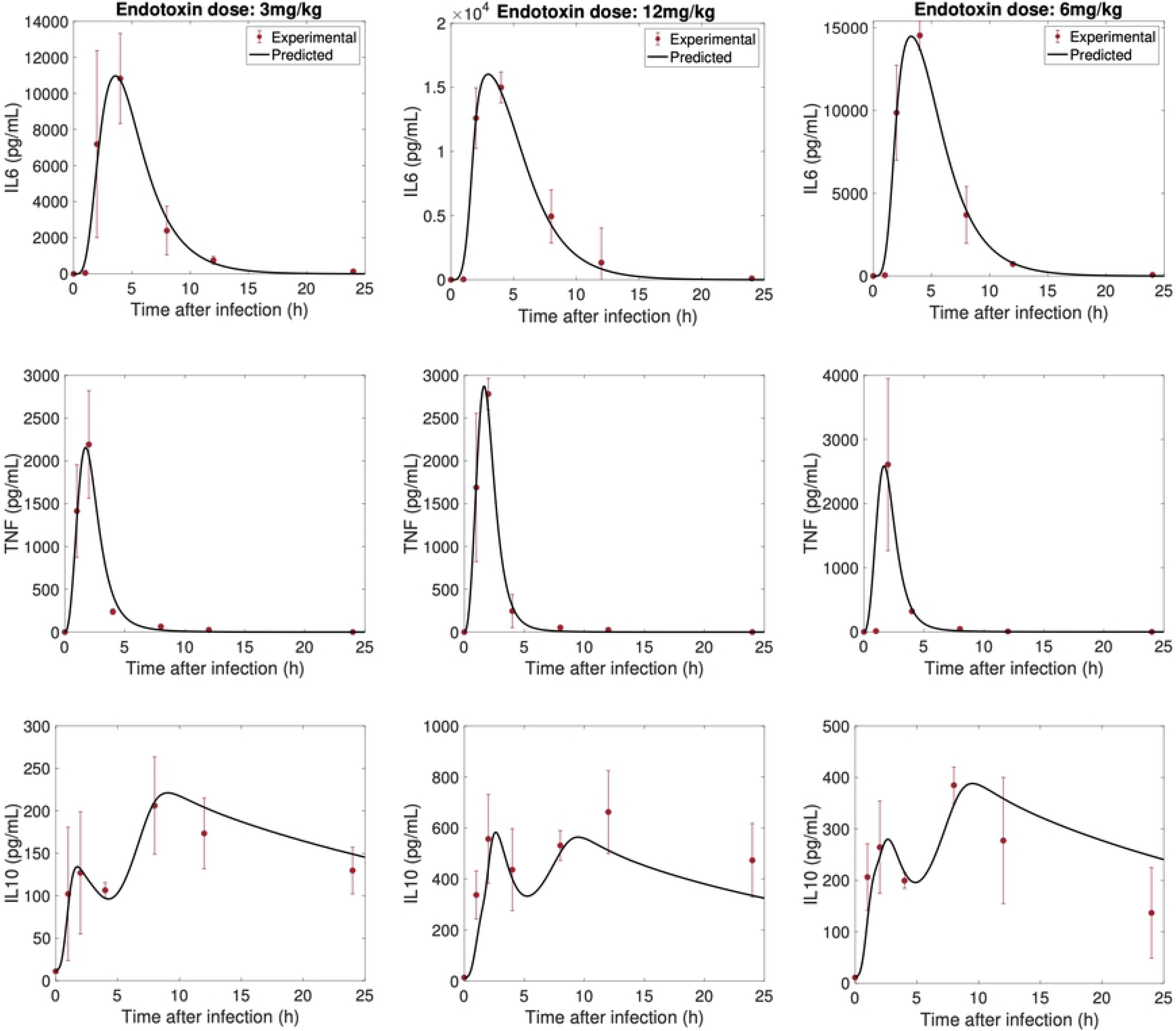
Predicted cytokine time profiles. Comparison of predicted time-courses of IL-6, TNF-*α* and IL-10 (solid line), against experimental data (circle) (mean ± SD), in response to endotoxin challenge at dosages of 3 mg/kg, 12mg/kg and 6mg/kg.

Model predictions were validated by comparing model simulations with available data at a 6 mg/kg endotoxin challenge level. These data were not used in the parameter fitting. The results are shown in Fig 4, right column. In general, model predictions of the measured cytokines are in good agreement with the experimental data. We observe that TNF-*α* concentration is overestimated at 1 h. This discrepancy may be explained by the apparent inconsistency between the data collected at 1 h with samples collected at the same time point for endotoxin dose levels of 3 and 12 mg/kg. A similar model behavior was observed in the initiating article by [33]. We note also that the model over-predicts IL-10 concentration after 10 h.

### Effect of infection timing on immune response

It is well established that the survival of mice after endotoxemic shock varies with the time of administration of bacterial insult. Studies have shown an increased lethality towards the end of the resting phase in controlled light-dark experiments [52, 70, 71]. Since an organism’s ability to fight off an infection depends in part on having a sufficiently large population of cytokines, we investigate how the timing of infection affects cytokine dynamics. Interestingly, a discrepancy can be discerned in post-infection IL-10 dynamics: some studies reported a single peak [91], whereas others reported two peaks [33]. We hypothesize that this discrepancy can be explained by the different infection timing (unfortunately infection timing was not reported in those studies).

We first seek to explain how the time of infection affects its lethality. Model simulations predict more tissue damage results from an infection administered at CT12 compared to CT0 (Fig. 5D). This difference can be explained in terms of the organism’s sensitivity to LPS, which is predicted to be highest at CT12, consistent with the observed phenotype of increased cytokine release [15, 52]. This increased sensitivity to LPS can be explained by a mismatch in the acrophases of REV-ERB and ROR in particular (see Fig 6). REV-ERB crests while ROR attains its minimum at CT12. Thus ROR inhibition of TNF-*α* and IL-6 is considerably reduced, hence allowing for a greater production of the cytokines. At the same time, the inhibition of IL-6 and IL-10 by REV-ERB is maximized, but since IL-10 also inihibits IL-6, its inhibition by REV-ERB repeals its action on IL-6, leading IL-6 to more than double (Fig 5A). We note that TNF-*α* increases less than IL-6 due to the inhibitory action of CRY which increases as REV-ERB decreases (see Fig 1 and Fig 6). The larger increase in cytokine populations following a CT12 infection results in more tissue damage, compared to a CT0 infection (Fig. 5).

**Fig 5.**
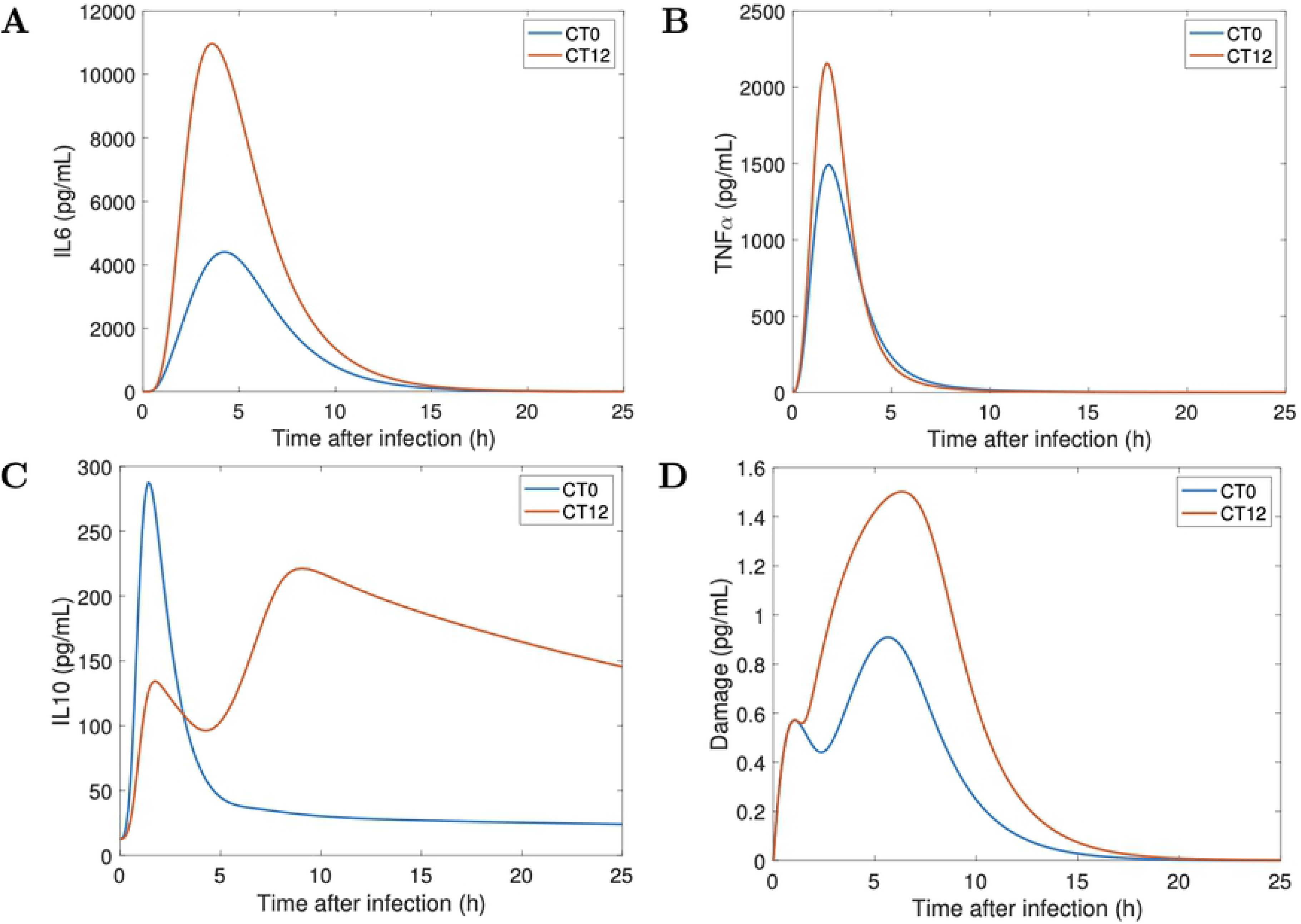
Inflammatory response after infection at CT0 and CT12. Model simulations of the time course of *IL-6*, *TNF-α*, *IL-10* and the damage marker for the control model in response to endotoxin dose 3mg/kg administered at CT0 and CT12

**Fig 6.**
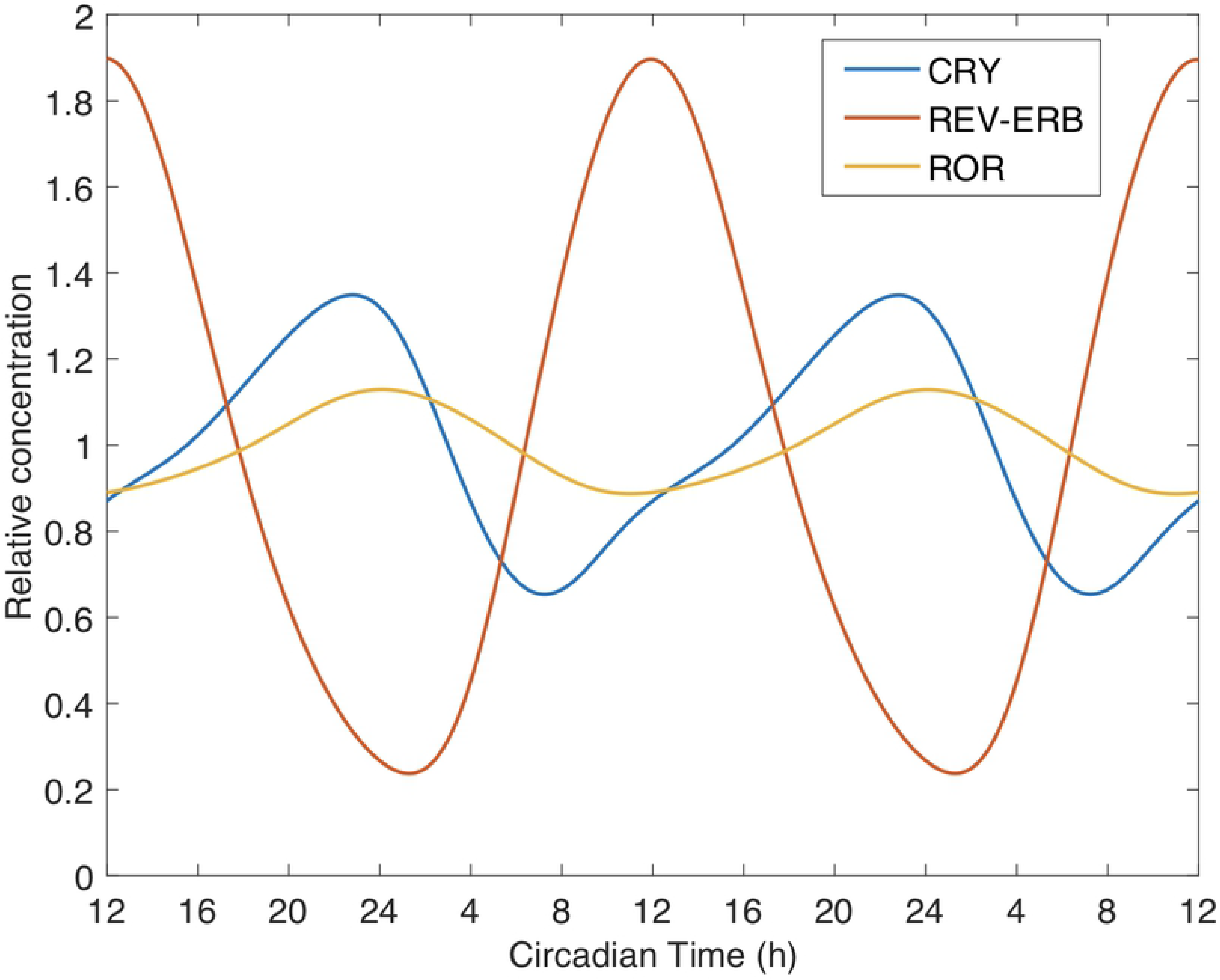
Main clock genes involved in the inflammatory response. Normalized temporal expression profiles of CRY, REV-ERB and ROR proteins, relative to their respective mean value.

It is noteworthy that our model predicts a single peak in the expression of the anti-inflammatory cytokine IL-10 when the host is infected at CT0 versus two peaks when the infection occurs at CT12 (Fig 5C). This behavior persists across different doses of endotoxin (results not shown), which indicates that the immune response might be different at CT0 compared to CT12 regardless of the extent of the infection. We hypothesize that the different timing of infection may explain the single peak versus two peaks in IL-10 time profiles in previous studies [33, 91]

A closer look at the dynamics of REV-ERB revealed the following: at CT0 and without infection, REV-ERB is close to its minimum. An attack by pathogens inhibits *Bmal1*, causing REV-ERB to drop below its nadir. This is followed by an almost complete loss of IL-10 inhibition by REV-ERB, and thus leads to a single IL-10 spike. The high levels of circulating IL-10 limit the production of proinflammatory cytokines (Figs 5A,B). On the contrary, REV-ERB is close to its maximum at CT12, and while an attack by pathogens inhibits its production, REV-ERB concentration remains sufficiently high for the first few hours. Therefore, REV-ERB still inhibits IL-10 during the early stages of inflammation and only a weak peak in IL-10 emerges. This inhibitory action is later counteracted by the accumulation of circulating IL-6 and TNF-*α* which upregulate IL-10 production, hence explaining the rise of a second peak in IL-10 when the infection occurs at CT12. About ten hours after the onset of inflammation, REV-ERB finally drops below its normal minimum levels. This is similar to the case at CT0, and leads to sustained elevated levels of IL-10 for a few hours after the elimination of IL-6 and TNF-*α*. We hypothesize that the lower production of pro-inflammatory cytokines at CT0 is due to high levels of IL-10, which spikes earlier during endotoxemia due to the loss of REV-ERB. The second peak in IL-10 production can be an indicator of a stronger inflammatory response as can be seen at CT12.

### Circadian disruption alters host immune response

It has been reported that a challenge of LPS in mice with a disrupted circadian clock leads to a stronger inflammatory response and increased mortality [73, 74]. Particularly, Castanon-Cervantes et al. observed a sustained reduction in *Bmal1* transcript following CJL [74]. This is similar to findings by Hadden et al. for female mice [12]. Below we conduct simulations to illustrate that *how* circadian disruption alters immune response depends on (i) the time of infection, and (ii) sex of the organism.

When endotoxin is administered at the onset of the rest phase (CT0), our model predicts an increased production of IL-6 and TNF-*α* in CJL males compared to controls (Fig 7A, B). Similar results were shown in experiments on CJL male mice [75] and CJL male rats [76] where the animals were injected with LPS during the early rest period. This stronger response is also observed in CJL females, and is due to the 8h-advance circadian disruption which resets their clock to the middle of the active phase. Although CJL females produced more IL-6 than controls, they remained relatively close to baseline compared to their male counterparts (Fig 7). Unlike males, the upregulation of CRY in CJL females decreases TNF-*α* and IL-6 production, while the downregulation of REV-ERB leads to increased levels of IL-10 and therefore more IL-10-induced IL-6 inhibition. In sum, CJL mice suffer more tissue damage from LPS administered at CT0, with CJL males more so than females (Fig 7D).

**Fig 7.**
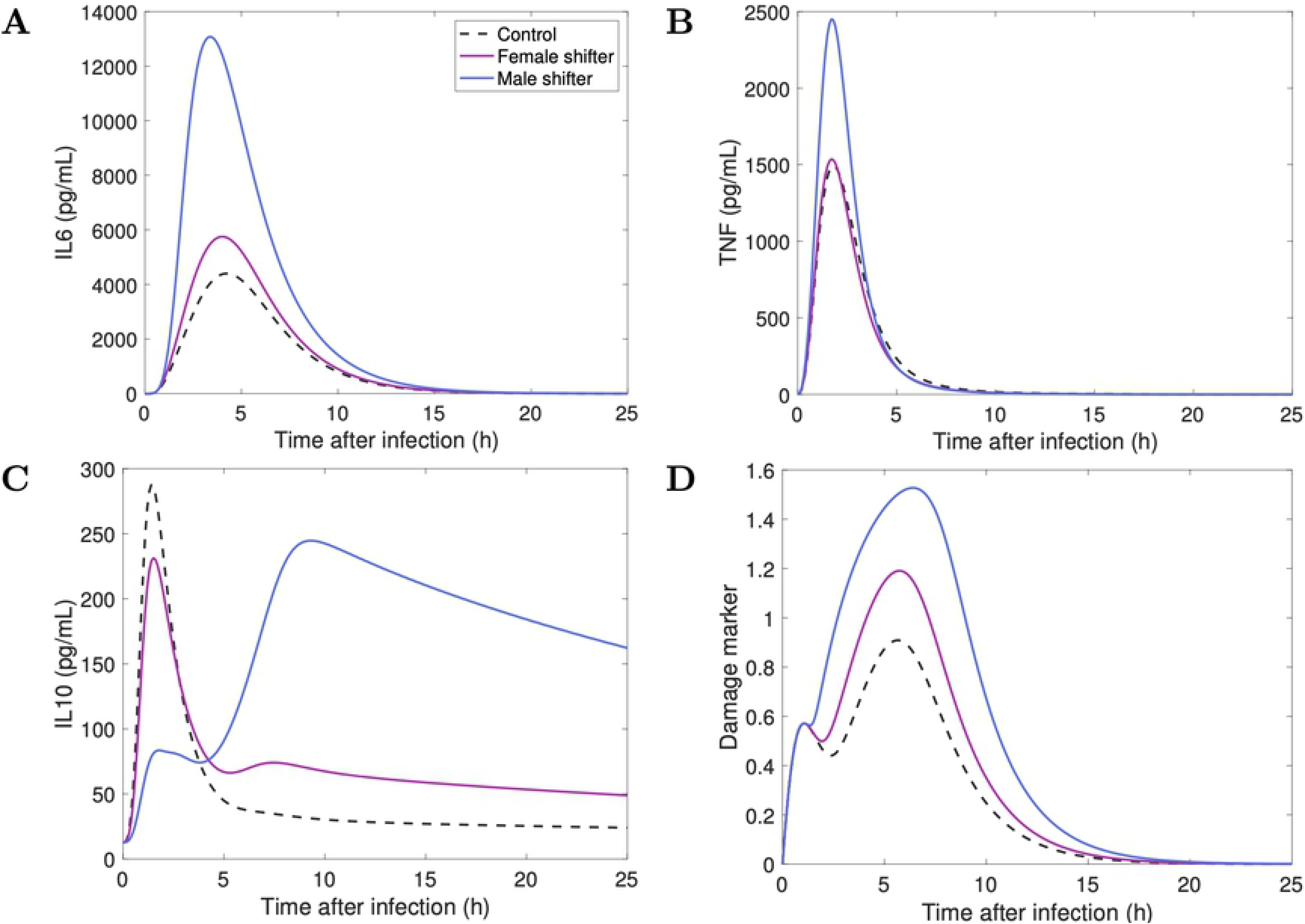
Sex-specific response to infection during CJL at CT0. Model simulations of the time course of *IL-6*, *TNF-α*, *IL-10* and the damage marker for controls (black) against CJL males (blue) and CJL females (pink) in response to endotoxin challenge of 3mg/kg administered at CT0.

The trend is reversed at CT12. Fig 8 shows that CJL mice have blunted IL-6 and TNF-*α* responses. We recall that REV-ERB is upregulated in CJL males compared to CJL females and controls (Fig 2D). Higher levels of REV-ERB further inhibit the production of IL-10, which then releases its inhibitory action on pro-inflammatory cytokines. This explains why CJL males, while having a blunted immune response, still produce more cytokines than CJL females. The downregulation of IL-6 following CJL has also been observed in all-male mice experiments [77–79]. Moreover, the weaker immune response in both CJL males and females is explained by their internal circadian disruption (8-h phase advance) which puts them at a circadian phase similar to CT4 in control mice. The reduced production of cytokine is represented by a lower damage marker and could indicate an inadequate and disrupted immune response (Fig 8D), particularly for CJL females. Our model predictions at CT12 are consistent with experimental reports that the immune system in females is detrimentally affected more than that of males during CJL [80].

**Fig 8.**
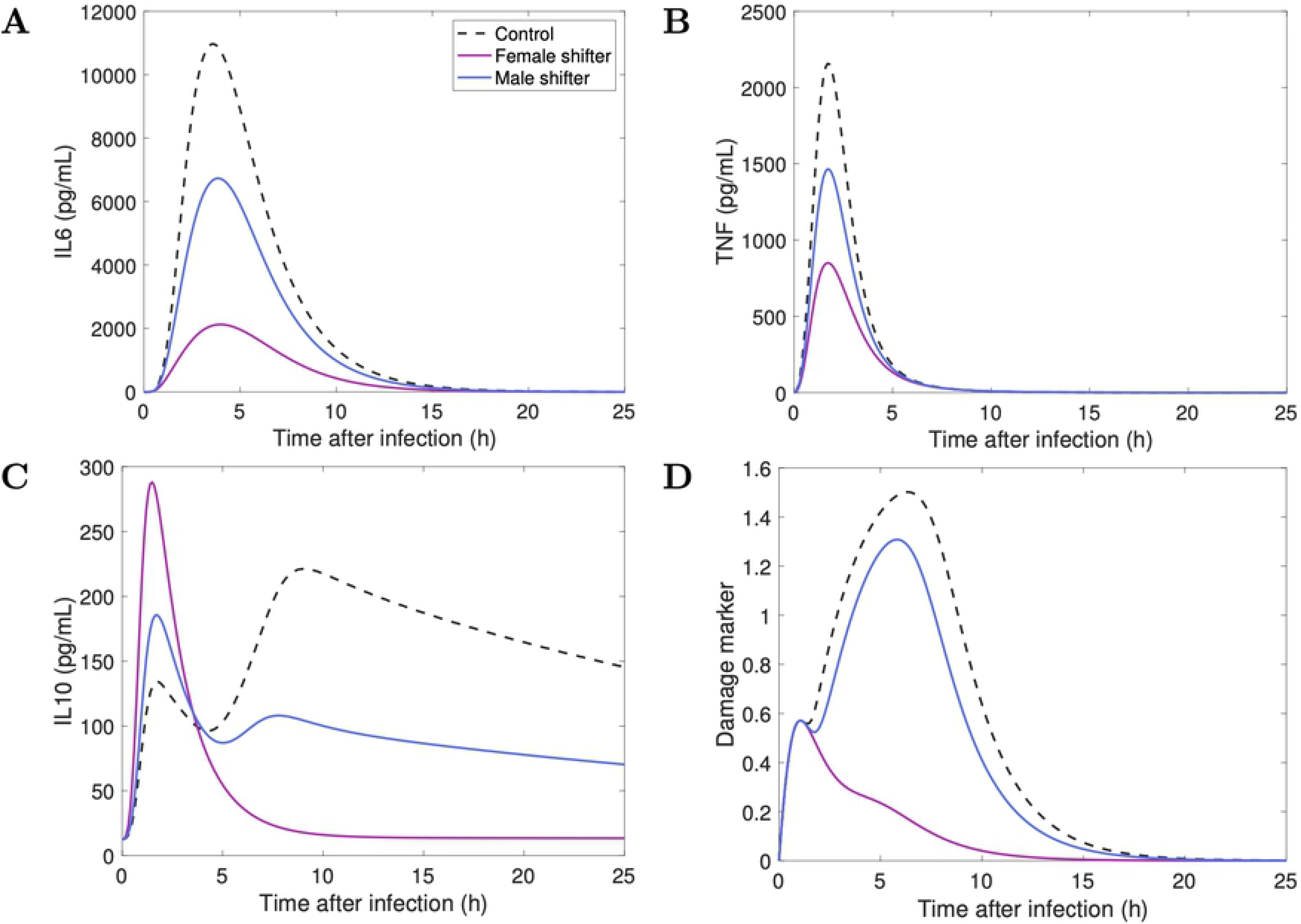
Sex-specific response to infection during CJL at CT12. Model simulations of the time course of *IL-6*, *TNF-α*, *IL-10* and the damage marker for controls (black) against CJL males (blue) and CJL females (pink) in response to endotoxin challenge of 3mg/kg administered at CT12.

### Effect of infection timing on immune response: Beyond CT0 and CT12

Previous studies have shown, in the absence of CJL, increased lethality towards the end of the resting phase, approximately 2 h before the onset of activity [70, 71]. We tested our baseline (no CJL) model predictions for different time points of infection, and indeed our results show higher sensitivity to LPS infection at CT9, followed closely by CT12 (see Fig 9A-D: left column).

**Fig 9.**
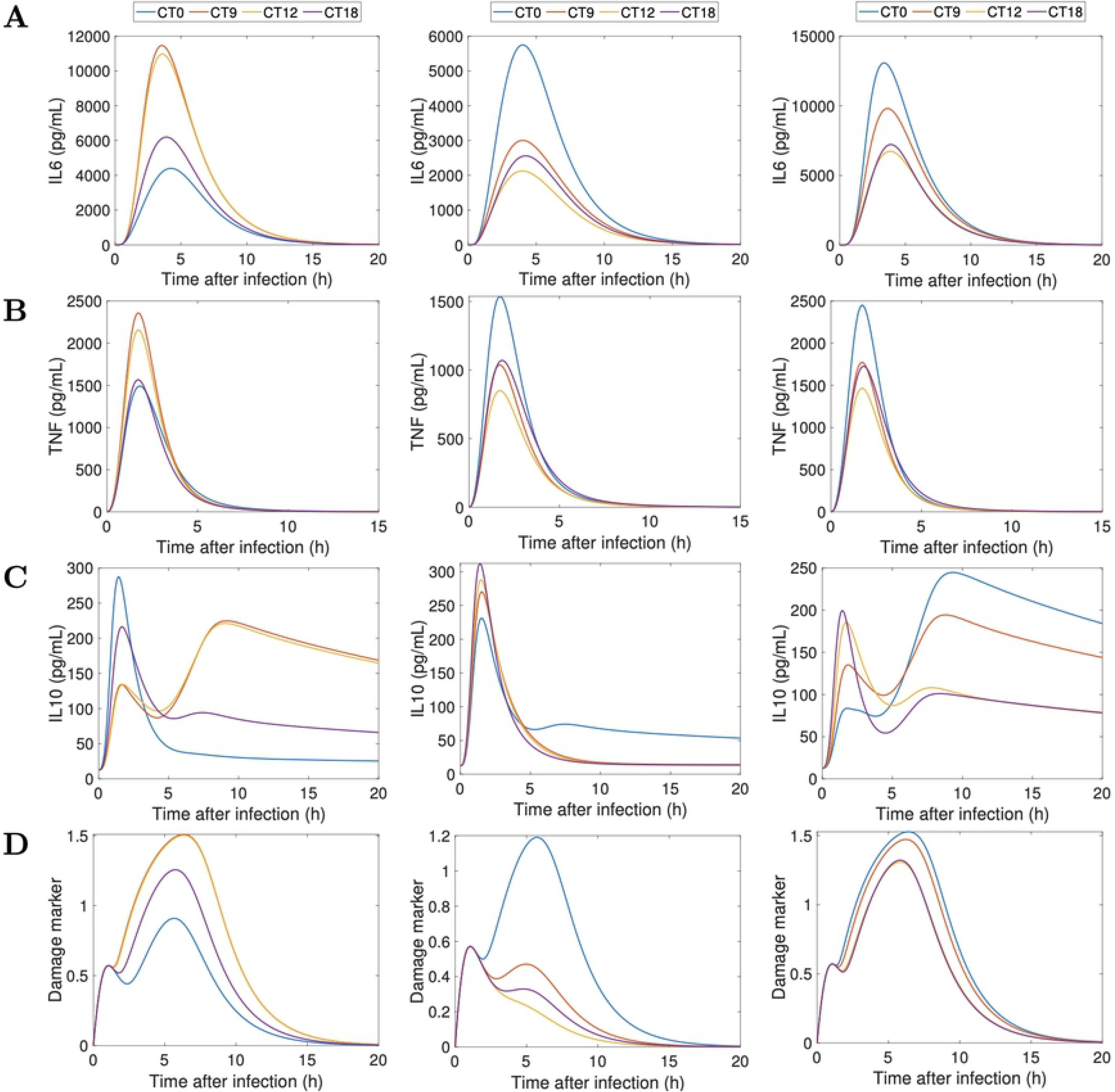
Simulated acute inflammation across different circadian times. *IL-6* (A), *TNF-α* (B), *IL-10* (C), *Damage marker* (D). The endotoxin dose is 3mg/kg. Left column, no CJL; middle column, CJL females; right column, CJL males.

We also conducted simulations under CJL. As shown in Fig 9A-D (middle and right columns), CJL males and females are more sensitive to LPS than controls at CT0, and exhibit reduced induction of cytokines at CT12. This supports our hypothesis that the CJL induced by an 8-h phase advance of the lung circadian clock would reverse the times of lowest and highest sensitivity to LPS. Interestingly, CT18 appears to be a time of lesser sensitivity to LPS regardless of the experience of the host: CJL or normal lighting conditions (Fig 9).

In general, the extent of sequelae experienced by male and female mice varied across the circadian day. Females had a more disrupted inflammatory response closer to CT12. Fig 9 (middle column) shows that CJL females were not able to recover the second peak in IL-10 at any of the times tested. This means that CJL females did not produce as much pro-inflammatory cytokines (e.g. CT9, CT12) as controls. Compared to CJL males, CJL females produce less cytokines overall during acute inflammation. Males, however, lost their circadian gating of cytokines closer to CT0 and produced greater amounts of pro-inflammatory cytokines, making them more susceptible to sepsis at that time [81, 82].

## Discussion

The circadian clock is responsible for the daily rhythms in immune functions [83]. A growing body of research supports the role of clock genes in regulating cytokines before and during infection [84]. In a reciprocal fashion, immune agents can impact the clock [85]. To investigate the interplay between the immune system and the circadian clock, we developed a mathematical model incorporating the bidirectional coupling between the lung circadian clock and the acute inflammatory response. We adapted this model to study the sexual-dimorphic effects of shift work (a.k.a. CJL) on both the clock mechanism and inflammation. It has been recognized that shift work has a negative impact on health [86] and a better understanding of the mechanisms by which disruption of circadian rhythms affects immunity, and how that effect differs between males and females, may help the development of chronotherapies for treating shift work-related disorders such as shift work sleep disorder (SWSD) and its related health-risks.

Given that sensitivity to LPS is highest at CT12 in mice [15, 52], we showed circadian disruption induced by an 8-h phase advance reduces cytokine production at this time while exacerbating the response at CT0. Remarkably, our models predict a second peak in the production of the anti-inflammatory cytokine IL-10 when the immune system is poised for attack (e.g. CT12). At times when the immune system is undergoing regeneration and repair (e.g. CT18-CT6) [15], the model predicts a single peak in IL-10 because the inflammatory response is weaker. The recurrence of this pattern at different doses of LPS implies the existence of qualitative differences at CT0 compared to CT12.

Overall, our results show that the extent of sequelae experienced by male and female mice depends on the time of infection. Females suffered more severe sequelae than males when infected during the late rest or early active periods. Specifically, IL-6 and TNF-*α* production at CT12 was greatly lowered in females. This response is not potent enough to maintain long term control of the infection. Female mice produce less cytokines overall during acute inflammation when compared to males. Nonetheless, males also suffer from circadian-induced immune disruption. Their higher levels of IL-6 and TNF*α* and damage could increase their susceptibility to sepsis.

The modulation of circadian activity by cytokines has been reported over the last years. For instance, TNF*α*-incubation has been shown to suppress *Per* gene expression *in vitro* and *in vivo* in mice [58] as well as *Cry1* [87]. Some recent work by [88] and [89] reveals that TNF*α* modulates the transcription of *Bmal1* through the up-regulation of *Rorα*. The present model does not include the direct links from cytokines to clock genes and proteins, but those links can be incorporated into future extensions of the model. We note that different lengths of phase advances and phase delays in the expression of clock genes could lead to different immune responses in males and females. These different scenarios provide an avenue for future research on immune-circadian regulation. The development of mathematical models that investigate the role of circadian rhythms in immunity and vice versa help our understanding of the dynamics involved in the interplay between these two systems.

In conclusion, our results suggest that circadian disruption due to shift work is primarily mediated by the circadian disruption of REV-ERB and CRY. REV-ERB in particular acts as an equilibrist by negatively affecting the expression of pro-inflammatory cytokine IL-6 and anti-inflammatory cytokine IL-10. We also showed the importance of sexual dimorphism in the magnitude of the inflammatory response during CJL. A functional and rhythmic clock confers immunoprotection and improves organismal fitness [90]. Thus, it is important to understand the molecular mechanisms that link the clock to immune functions, particularly during unrest caused by behavioral changes such as shift work.

## Supporting information

**S1 File. Supplemental tables.** (Table S1) List of variables, (Table S2) list of the differential equations defining the mathematical model; relates to Fig 1. (Tables S3-S10) Nominal parameter set of the mathematical model.

## Acknowledgments

This research was supported by the Canada 150 Research Chair program and the NSERC Discovery award.

